# Starvation reduces thermal limits of the widespread copepod *Acartia tonsa*

**DOI:** 10.1101/2023.06.20.545723

**Authors:** Gaia A. Rueda Moreno, Matthew C. Sasaki

**Affiliations:** Department of Biology, New York University; Department of Marine Sciences, University of Connecticut

**Keywords:** copepod, temperature, thermal limit, climate change, starvation

## Abstract

Organismal thermal limits affect a wide range of biogeographical and ecological processes. Copepods are some of the most abundant animals on the planet, and play key roles in aquatic habitats. Despite their abundance and ecological importance, there is limited data on the factors that affect copepod thermal limits, impeding our ability to predict how aquatic ecosystems will be affected by anthropogenic climate change. In a warming ocean, one factor that may have particularly important effects on thermal limits is the availability of food. A recently proposed feedback loop known as “metabolic meltdown” suggests that starvation and exposure to high temperatures interact to drastically reduce organismal thermal limits, increasing vulnerability to warming. To investigate one component of this feedback loop, we examined how starvation affects thermal limits (critical thermal maxima: CTmax) of *Acartia tonsa*, a widespread estuarine copepod. We found that there was no effect of short duration exposure to starvation (up to two days). However, after three days, there was a significant decrease in the CTmax of starved copepods relative to the fed controls. Our results provide empirical evidence that extended periods of starvation reduce thermal limits, potentially initiating “metabolic meltdown” in this key species of coastal copepod. This suggests that changes in food availability may increase vulnerability of copepods to increasing temperatures, amplifying the effects of climate change on coastal systems.

## Introduction

The acquisition of nutrition is a fundamental challenge for organisms. Environmental conditions have well-known effects on the ability to find, capture, and ingest food via a direct influence on organismal performance. Temperature, for example, has strong effects on metabolic rates in ectothermic organisms, linking both performance and energetic requirements to the thermal environment (Brown *et al*., 2004; Johnston *et al*., 2022). As energetic requirements tend to increase with warming, we might expect to see increasing consumption pressure on autotrophs and lower trophic levels as ocean warming proceeds, with consequences for community structure and the distribution of biomass (Archibald *et al*., 2022). Nutrition may also affect how environmental conditions impact organisms, however, by modifying the underlying sensitivity to changes in the experienced environment (Huey & Kingsolver, 2019; Litchman & Thomas, 2023). As anthropogenic climate change is driving long-term ocean warming and increased frequency of disturbances like marine heat waves (Hobday *et al*., 2016; Oliver *et al*., 2018; Smale *et al*., 2019; Harvey *et al*., 2022; Johnston *et al*., 2022; Li & Donner, 2022), these feedbacks between feeding and sensitivity to environmental conditions are crucial to consider.

The interactions between starvation and upper thermal limits are still unknown for many taxa. Most studies have focused on terrestrial arthropods, examining species in Diptera (Kalra *et al*., 2017; Mitchell *et al*., 2017; Gotcha *et al*., 2018; Manenti *et al*., 2018), Coleoptera (Scharf *et al*., 2016; Chidawanyika *et al*., 2017), Hymenoptera (Nguyen *et al*., 2017; Gonzalez *et al*., 2022), Hempitera (DeVries *et al*., 2016), and Lepidotera (Mir & Qamar, 2018; Mutamiswa *et al*., 2018). There is strong variation in the effects of starvation on thermal limits across this body of work. The duration of starvation is likely important to consider, with little to no effect of shorter periods of starvation (DeVries *et al*., 2016; Gonzalez *et al*., 2022) and decreases in thermal limits after longer periods (Nguyen *et al*., 2017). However, even relatively short exposures have been observed to reduce thermal limits in some taxa (Manenti *et al*., 2018; Mir & Qamar, 2018). Further, a number of studies have reported that starvation actually increases thermal limits (Bubliy *et al*., 2011; Kalra *et al*., 2017; Gotcha *et al*., 2018), possibly due to starvation-induced changes in energy allocation. Within aquatic ectotherms, experiments on the freshwater amphipod *Gammarus fossarum* revealed that starvation improved survival during acute heat stress (Semsar-kazerouni *et al*., 2020). This improvement in thermal limits is not observed in fish or octopus systems (Lee *et al*., 2016; Uriarte *et al*., 2018).

The species-specific nature of starvation effects on thermal limits is an important observation for predictions about the response of communities to climate change. Amplifying the direct effects of starvation on thermal limits, Huey & Kingsolver (2019) describes a process termed ‘metabolic meltdown’, in which exposure to high temperatures, reduced food intake, and decreasing thermal limits act synergistically to drastically decrease tolerance to a warmer environment. Since a key component of metabolic meltdown is the reduction of thermal limits under reduced food intake, variation in starvation effects across taxa may determine the relative vulnerability of community members to events like heatwaves, thus shaping community dynamics in a changing climate.

Copepods are the some of the most abundant animals on the planet, and dominate planktonic communities in the coastal ocean (Turner, 2004). By nature of their abundance this group plays key ecological and biogeochemical roles in aquatic systems (Steinberg & Landry, 2017; Brun *et al*., 2019; Pinti *et al*., 2023). In particular, copepods are important consumers of primary productivity, and act as a crucial linkage between phytoplankton and higher trophic levels (Castonguay *et al*., 2008). Climate change therefore has the potential to impact aquatic ecosystem functions as well as human fishery systems directly through effects on copepod populations. However, despite their abundance and ecological importance, there is limited data on environmental control of copepod thermal limits, including how their thermal limits are affected by starvation. This impedes our ability to predict how copepod populations may be affected by co-occurring changes in temperature and food availability over both short (e.g. - seasonal changes) and long timescales (e.g. - anthropogenic climate change). In this study we tracked changes in critical thermal maxima (CT_max_) in the widespread copepod *Acartia tonsa* during extended starvation to test the hypothesis that food deprivation reduces thermal limits.

## Methods

### Copepod Cultures

The copepods used in this study were collected in July 2020 from Esker Point, Connecticut (41.3206N, -72.002W) by surface tow with a 63-um mesh net and solid cod end. Mature *Acartia tonsa* females and males were isolated from the tow contents, and used to initiate a laboratory culture which was maintained in 12 qt buckets. Cultures were kept in an environmental chamber at 18°C with a 12:12 light:dark cycle. A small aquarium pump ensured constant aeration. Copepods were fed *ad libitum* a mixture of three phytoplankton cultured in F/2 media under the same environmental conditions: a green flagellate, *Tetraselmis sp*.; a cryptomonad *Rhodomonas sp*.; and a small diatom, *Thalassiosira weissflogii*. This diet is regularly used to maintain large, active laboratory cultures of *A. tonsa* (Sasaki & Dam, 2021b).

### Measuring Thermal Limits

We used a custom set up to measure critical thermal maxima (CT_max_) of individual copepods, as described in (Sasaki *et al*., 2023). Briefly, this set up includes a reservoir, water bath, and temperature logger. The reservoir (a five gallon bucket) holds ∼15L of water, along with a 300-watt fixed output titanium water heater and two aquarium pumps. One pump vigorously circulates water within the reservoir while the other pumps water up into the water bath, a plexiglass tank that sits atop the reservoir. The water bath contains a series of test tube holders, used to position the experimental vessels (50 ml flat-bottom glass tubes) during the assay. When the pump is turned on, water floods the bath and then spills over back into the reservoir. In this arrangement, temperatures in the experimental vessels are slowly increased at a rate of between 0.1-0.3°C per minute, following temperatures in the reservoir. The final component is a small Arduino logger, which records temperature from three sensors placed inside tubes distributed throughout the water bath.

At the beginning of each CT_max_ trial, the water in the reservoir was adjusted to 18°C. The experimental vessels were filled with 10 mL of 0.2 um filtered sea water before being placed in the water bath, which when flooded brought the experimental vessels to the correct temperature as well. Individual copepods were placed into the vials (n = 10 per assay), and let acclimate for ten minutes at constant temperature. All copepods were checked during this time period for normal behavior. Individuals exhibiting abnormal behaviors were excluded from further analysis. After this resting phase, the water heater was turned on, initiating the temperature ramp. Simultaneously, the temperature logger began to record temperature and a stop watch began recording the time passed. Individuals were continuously monitored as water temperature increased. CT_max_ is generally defined as the temperature at which an individual ceases to respond to physical stimuli (Cowles & Bogert, 1944), indicating the onset of “ecological death” (the inability to escape lethal temperatures, predators, etc.). In *A. tonsa* this is indicated by cessation of movement, a lack of response to gentle physical stimuli (e.g. - slow flushing of the water in the tube with a transfer pipette), and abnormal body configuration (specifically - antennules pressed against the sides of prosome and a distinct dorsal tilt of the urosome). The time at which an individual began to exhibit these characteristics was recorded and then that individual’s experimental vessel removed from the water bath. After all individuals reached their CT_max_, copepods were photographed using a camera attached to an inverted microscope, and body size estimated using a scale micrometer and the software ImageJ (Schneider *et al*., 2012).

The times recorded during the trials were converted to CT_max_ values in degrees C using the temperatures recorded on the Arduino logging system. As there are between 1-10 vials being monitored at any point in the trial, the time at which an individual was recorded as having stopped responding to stimulus corresponds with the latest time (and therefore highest temperature) it could have reached its CT_max_. The period of time during which an individual could have reached its CT_max_ extends from this definite end point to the last time the individual was checked. As it generally takes around 5 seconds to check whether an individual has stopped responding, the duration of this period of time was estimated for each individual as the number of vials remaining in the water bath multiplied by 5 seconds. As a result, this uncertainty window decreased in length as the trial went on, until, for the final individual, the window includes just the amount of time it took to check whether the individual had stopped responding. For each individual, CT_max_ is estimated as the average temperature recorded by all three temperature sensors throughout the uncertainty window. We used this time-based method instead of directly monitoring the temperatures because i) it was more efficient to record the time than the temperature readings from three separate sensors, and ii) to reduce any sub-conscious bias stemming from past knowledge or expectations about the copepod thermal limits.

### Experimental Design

We used five replicate experiments to test our hypothesis that starvation reduces copepod thermal limits over time. Each experiment involved measuring a baseline CT_max_ for the culture, and then making daily CT_max_ measurements for two groups of copepods, a fed control group and a starved treatment group. To initiate each experiment, 10 mature females were isolated from the laboratory culture and maintained for 24 hours in 200 mL of an *ad libitum* food solution (800 ug C / L of *Tetraselmis*). Preliminary work showed that short exposure to three different prey options (*Tetraselmis sp*., *Rhodomonas sp*., and *Oxyrrhis marina*, a heterotrophic dinoflagellate) did not affect copepod thermal limits (Figure 1). After 24 hours, CT_max_ was measured as described above. This provided an initial baseline value for each experiment.

**Figure 1:**
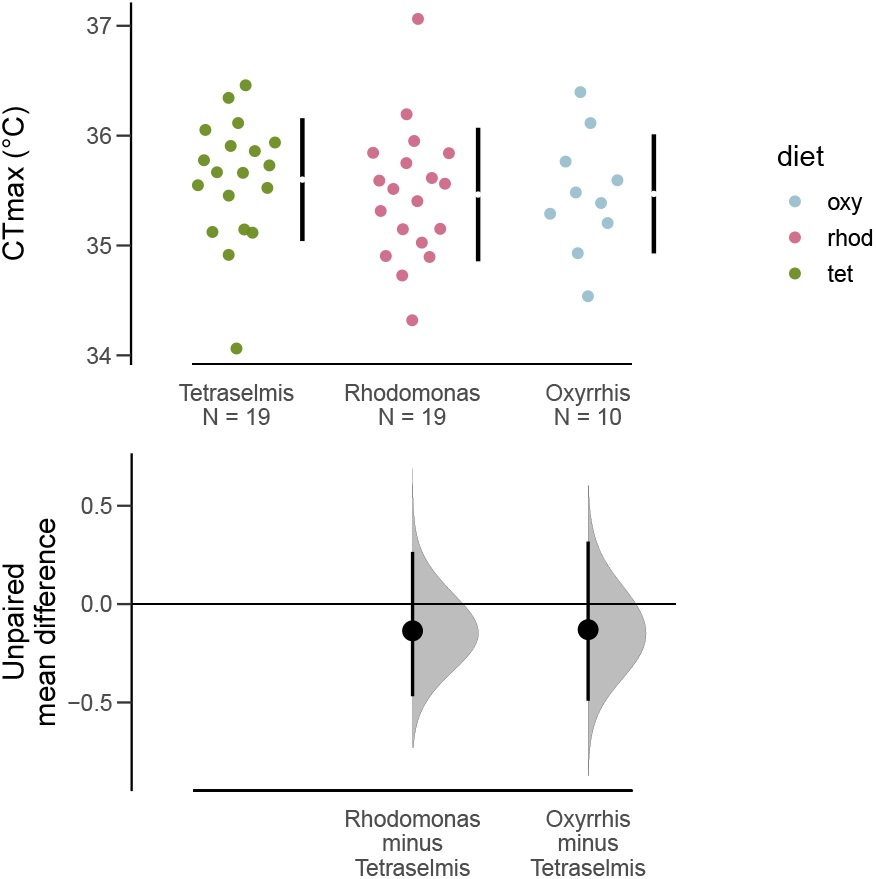
Estimation plots showing the effect of different diet treatments on thermal limits. Measurements were taken after 24 hours, and show that thermal limits were not altered by short exposure to the different diets.

On the same day as the baseline CT_max_ measurements, ∼90 mature females were isolated from the culture and divided into six groups. The six groups were randomly assigned to either the starvation treatment or the fed control treatment. The starvation groups were maintained in 0.2-um filtered sea water. The control group was provided with the food solution used prior to measuring baseline CT_max_ (800 ug C / L of *Tetraselmis*). Each group was kept in a 100 mL cup with a plastic cylinder nested within. The bottom end of the cylinder was covered by a 150-um mesh screen. Similar set-ups are often used to prevent egg cannibalization during egg production assays, as eggs of *A. tonsa* sink through the mesh to the base of the cup (Plough *et al*., 2018). In our case, this prevented females in the starvation group from acquiring nutrition via egg cannibalism. All groups were transferred to fresh media (either filtered sea water or food solution) on a daily basis throughout the experiment by gently removing the meshed column and placing it into a new 100 mL cup.

Copepods from these six groups were then used to measure thermal limits each day for five days, starting 24 hours after the females were first isolated. Each day, we measured CT_max_ values for ten copepods, selected at random from the six groups (see the accompanying code for the randomization script used). By repeating these measurements over the five day period, we were able to examine how the effects of starvation on CT_max_ changed as the duration of exposure increased. The second replicate experiment was ended after day three when all individuals from the starvation treatment died. Individuals from replicate experiment five were not photographed after the CT_max_ assays due to a malfunction with the imaging software.

### Statistical Analysis

All statistical analyses were performed with R (Version 4.1.3; R Core Team 2022). We examined the effects of starvation using mean difference as an effect size estimate. Confidence intervals for this effect size were estimated using non-parametric bootstrapping (Ho *et al*., 2019). Two sets of effect sizes were estimated. First, we examined the difference between the initial baseline CT_max_ values and CT_max_ values for fed and starved individuals during the experiment (e.g. - the difference between fed CT_max_ and the baseline CT_max_ on each of days one through five). The second effect size was the difference between fed and starved individuals on each day within the experiment. If CT_max_ does not change over time in the fed control group, these two effect sizes will provide similar results.

## Results

A total of 254 CT_max_ measurements were made across the five replicate experiments (153 from the fed controls and 101 from the starved treatment). Note that the difference between the number of fed and starved individuals reflects the five sets of initial baseline CTmax values measured (all experienced ‘fed’ conditions). The custom setup produced consistent ramping rates across assays (Figure 2). Ramping rates did however decrease over time in a consistent way within each assay due to the imperfect insulation of the bucket reservoir. Nonetheless, ramping rates were always between the target ramping rates of 0.1 - 0.3°C per minute, which have been used previously to measure CT_max_ for copepods (Jiang *et al*., 2009; Harada & Burton, 2019; Sasaki & Dam, 2021a).

**Figure 2:**
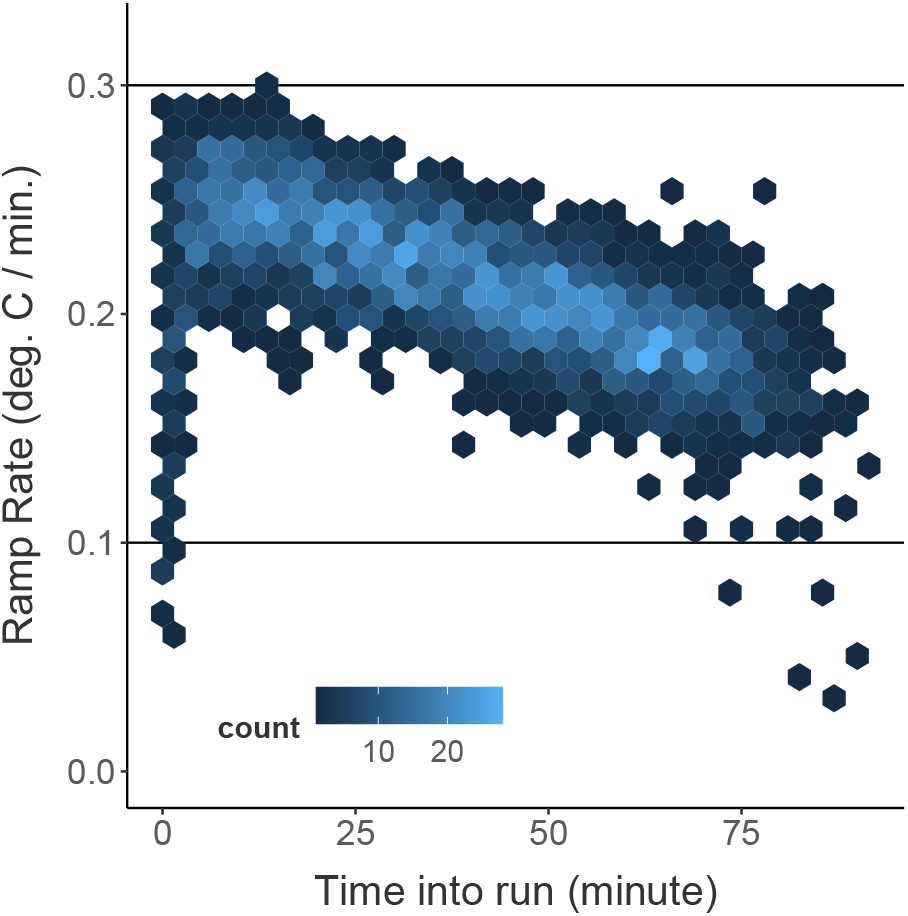
A heatmap of the observed ramping rates (the increase in temperature per minute) during the CTmax trials. The plane is divided into regular hexagons, which are shaded according the frequency of the encompassed ramping rates. Ramping rates were always between the target values of 0.1 and 0.3°C per minute.

Thermal limits of the fed control individuals were consistent with the baseline CT_max_ values across the entire experiment. We will therefore focus on the direct comparisons between fed controls and starved individuals. Mean CT_max_ in the starved group gradually decreased over the course of the five day starvation period relative to the fed controls (Figure 3). There was no difference between thermal limits of fed and starved individuals on days 1 or 2. Thermal limits were approximately 1°C lower by day 3 and continued to decrease by approximately 2°C per day after that. By day 5, starved individuals had thermal limits that were approximately 5°C lower than those of control individuals. There was, however, also a large increase in the variance of individual thermal limits in the starved treatment; several individuals maintained thermal limits similar to those observed in the control individuals while others had thermal limits as low as 22°C. There was no significant effect of body size on CT_max_ during any of the experimental days (Figure 4).

**Figure 3:**
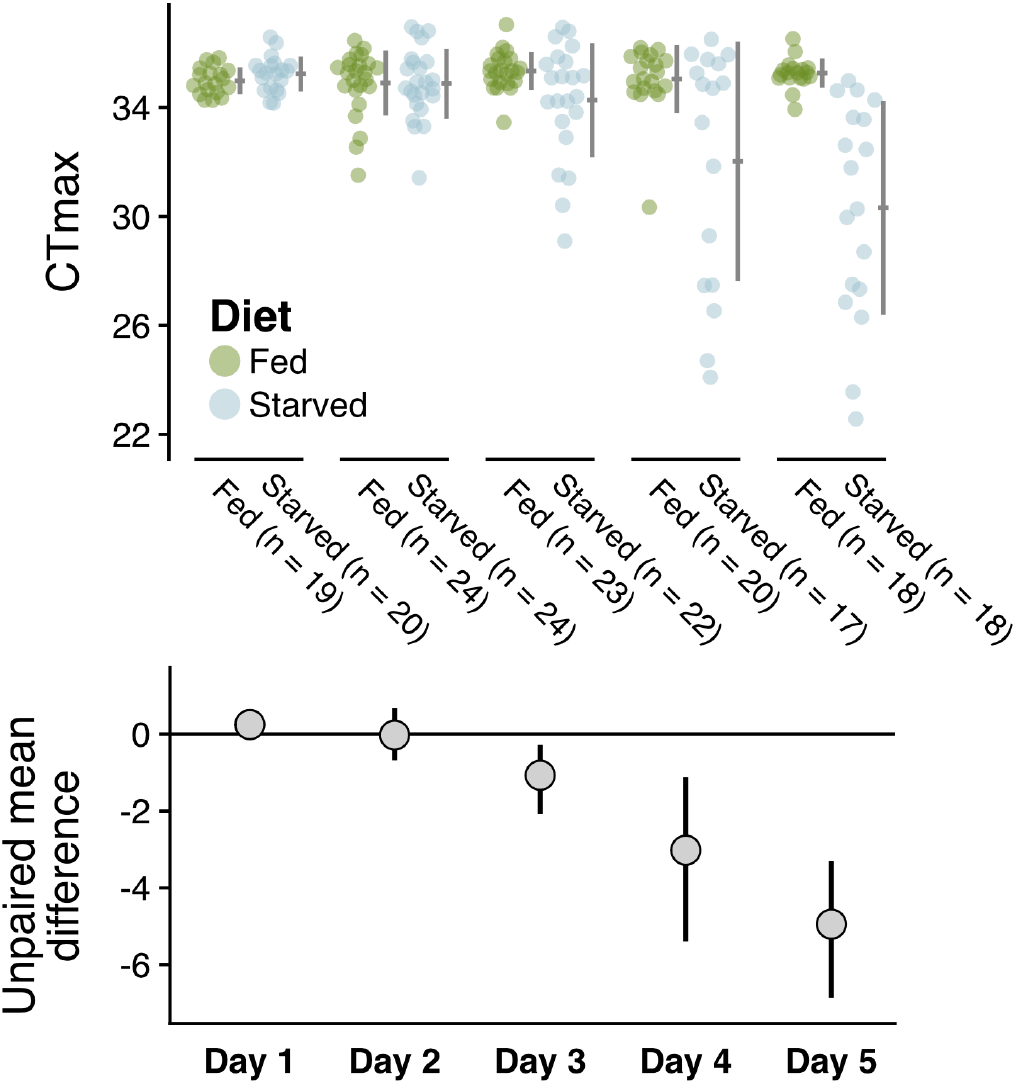
Estimation plots depicting the gradual reduction in thermal limits (measured as CTmax) in the starvation group relative to the fed control group. The top panel shows the raw CTmax values, with fed and starved individuals in green and blue, respectively. The bottom panel shows the calculated effect sizes for each comparison, along with 95 percent confidence intervals.

**Figure 4:**
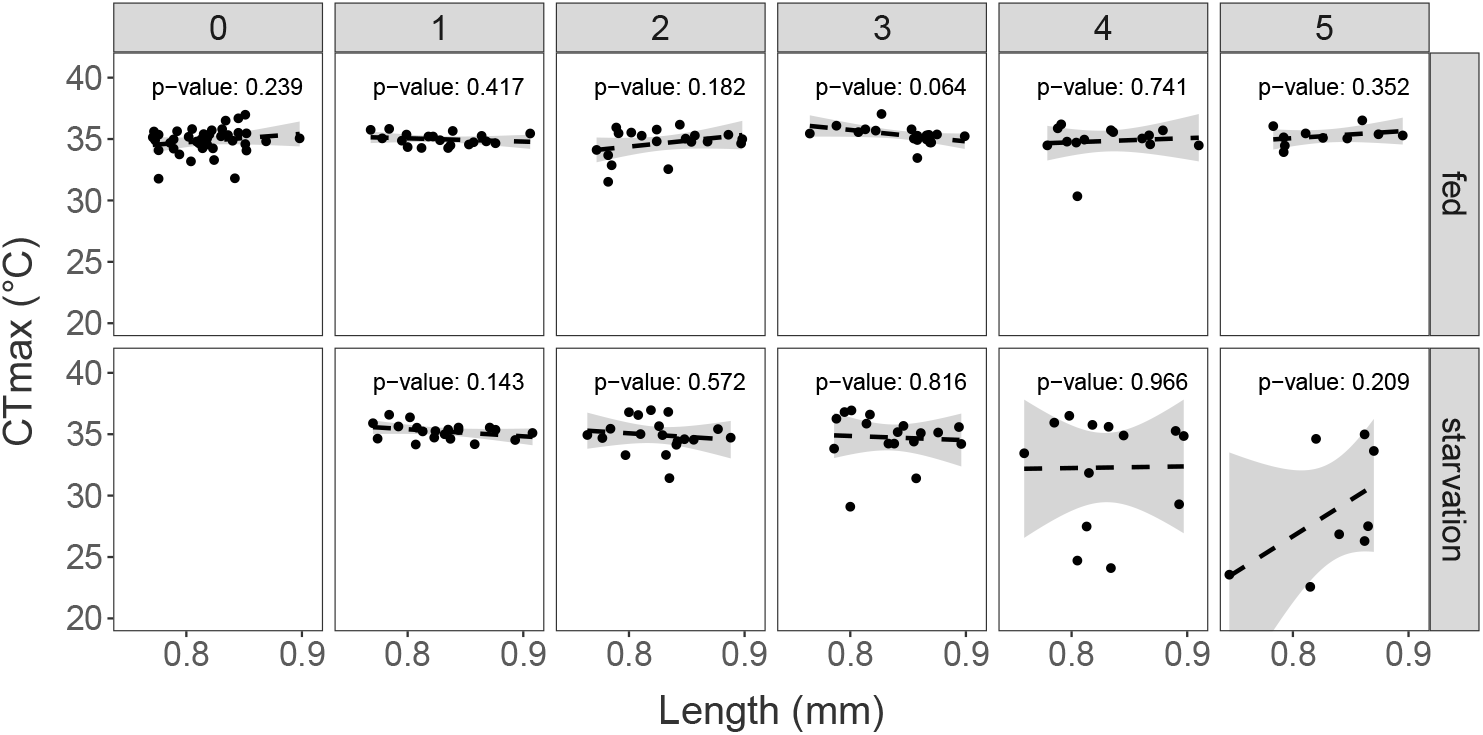
Thermal limits (CTmax) plotted against individual body size for each day, separated by treatment. A linear regression is shown, along with the associated p-value for each day. Thermal limits are not correlated with body size on any of the experimental days.

## Discussion

Marine biota are threatened by both ongoing warming and changes in food quality and quantity. Examining the interactions between starvation and environmental sensitivity may therefore provide useful insights about the responses of marine communities to climate change. We tested the hypothesis that food deprivation reduces thermal limits in the important coastal copepod species *Acartia tonsa*. Using a series of replicated laboratory experiments, we show that the upper thermal limits of *Acartia tonsa* are substantially reduced after just a few days without food, although this reduction was highly variable across individuals.

Previous studies have shown a wide range of starvation effects on upper thermal limits. The range of responses implicates several potential mechanistic bases starvation effects on thermal limits. In systems where thermal limits increase due to starvation, enhanced lipid accumulation at the initiation of starvation and changes in energetic allocation have been suggested as a mechanistic basis (Djawdan *et al*., 1998; Sokolova, 2013; Klepsatel *et al*., 2016). Here, however, we observed a decrease in thermal limits resulting from starvation. In other systems where this has been observed, the depletion of energetic reserves and the accompanying reduction in metabolic rates are often suggested as a potential physiological basis. Starvation tolerance in copepods may be mediated by similar processes. While Acartiid copepods have relatively small energetic reserves compared to other calanoids (e.g. the large lipid reserves of *Calanus* species), these energetic reserves may still buffer against deleterious effects of starvation over several days (Hirahara & Toda, 2018). The 3-day buffer against decreases in thermal limits that we observed in this study contrasts the rapid drop-off of egg production and respiration rates in starved *A. tonsa* females (Parrish & Wilson, 1978; Kiørboe *et al*., 1985; Thor, 2003). This may indicate different biochemical bases, or that environmental tolerance and survival is prioritized in energetic allocation. It is important to address the physiological basis for the observed patterns and disentangle cause and consequence (MacMillan, 2019). One approach may be to examine how patterns in respiration rates and heat shock protein expression change during starvation and heat stress (both individually and when combined).

Developing an understanding of how changing environmental factors affect the thermal performance of copepods is crucial due to their role as the linkage between primary producers and larger consumers in aquatic foodwebs. Moreover, the factors affecting the physiological limits of *A. tonsa* specifically are important to understand given the key role this widely distributed species plays in coastal communities (Turner, 1981, 2004). It is well known that phenotypic plasticity and genetic differentiation both affect the upper thermal limits in this species (Sasaki & Dam, 2019), potentially facilitating its wide distribution. Factors that limit physiological performance in this species are also important to understanding for predicting invasion risk across temperate and subtropical estuaries, where anthropogenic introduction of this species may alter community dynamics (Aravena *et al*., 2009; Svetlichny & Hubareva, 2014). Additional experiments are needed to understand how starvation effects on upper thermal limit may change across temperature gradients. However, assuming an energetic basis to the reduction in thermal limits observed, we might expect to see larger, more rapid decreases in thermal limits when high temperatures and starvation co-occur, potentially inhibiting invasion of warm, oligotrophic regions.

Another open question is to what extent sex-specific differences in starvation sensitivity or developmental effects of food limitation may influence population dynamics. Females in this species are known to be both more resistant to starvation (Finiguerra *et al*., 2013) and to have higher thermal limits than males(Sasaki *et al*., 2019). Increased male sensitivity to starvation relative to females (resulting in a more rapid or a larger decrease in thermal limits) may amplify negative population demographic effects of co-occuring food limitation and exposure to high temperatures. The effects of developing at different food levels may also influence these dynamics. Developmental patterns may be strongly influenced by food limitation, with increases in total development time, and decreases in adult size (Vidal, 1980; Hirst *et al*., 2003). This may dampen starvation effects on thermal limits simply because smaller adults typically require less food to maintain basal energetic demands. This may result in an increased capacity to maintain thermal limits under starvation in individuals that completed development at low food levels. While ontogenetic patterns in respiration rates are resilient to heat stress in *Acartia tonsa* (Holmes-Hackerd *et al*., 2023), the effects of food limitation during development may have carry-over effects on adult respiration and energetic requirements that may modify these predictions. These carry-over effects are, however, still unknown.

Regardless of the mechanistic basis, our results highlight the importance of understanding temporal scales of physiological variation for predictions about vulnerability. Over short timescales, *Acartia tonsa* females are able to maintain thermal limits under starvation. This suggests that starvation is unlikely to increase vulnerability of natural populations to acute events like heatwaves. Longer timescale decreases in food availability, however, may have dire consequences for populations in a warming ocean. Although research on large-scale changes in phytoplankton biomass due to climate change lacks consensus (Boyce *et al*., 2010; Taucher & Oschlies, 2011; Archibald *et al*., 2022), broad decreases in vertical mixing associated with the intensified thermal stratification and shifts in the phenology of the spring bloom to earlier in the year (Sommer & Lengfellner, 2008) may drive an increase in small, motile phytoplankter and heterotrophic microzooplankton in many regions (Winder & Sommer, 2012). Primarily herbivorous copepods may therefore be particularly vulnerable to warming compared to omnivorous species like *Acartia tonsa*, which may avoid starvation and the accompanying reduction in thermal limits by feeding on heterotrophic microzooplankton. Given the large contributions of herbivorous species to carbon flux in marine systems (Pinti *et al*., 2023), these shifts in community composition driven by differential environmental sensitivity would have important biogeochemical consequences.

